# Bone Marrow- and Umbilical Cord-Derived Mesenchymal Stem Cell Secretome Alters Gene Expression and Upregulates Motility of Human Endometrial Stromal Cells

**DOI:** 10.1101/2022.11.12.516251

**Authors:** Qingshi Zhao, Karla Larios, Yahaira Naaldijk, Lauren Sherman, Anat Chemerinski, Kennisha Okereke, Pranela Rameshwar, Alexander Lemenze, Nataki C. Douglas, Sara S. Morelli

## Abstract

**Introduction:** Cyclic regeneration of the endometrium, and its repair after parturition or injury, are crucial for successful reproduction. Mesenchymal stem cells (MSCs) derived from bone marrow (BM-MSC) and umbilical cord (UC-MSC) facilitate tissue repair via their secretome, which contains growth factors and cytokines that promote wound healing. Despite the implication of MSCs in endometrial regeneration and repair, the mechanisms remain unclear. This study tested the hypothesis that the secretome of MSCs from human BM and UC upregulates human endometrial stromal cell (HESC) proliferation, migration and invasion, and activates pathways to increase HESC motility.

**Methods:** MSCs were purchased from ATCC (BM-MSC-1) and cultured from the BM aspirate of three healthy female donors (BM-MSC-2-4), and from umbilical cords of two healthy male term infants (UC-MSC-1-2). Indirect co-culture of MSCs and hTERT-immortalized HESCs via a transwell system studied the effect of the BM-MSC and UC-MSC secretome on HESC proliferation, migration, and invasion. To study the effect of the MSC secretome on HESC gene expression, HESCs were exposed to the BM-MSC secretome via indirect co-culture for 24 h. Total RNA was extracted from HESCs for RNA sequencing (RNA-Seq). Differentially expressed genes (DEG) and significantly altered pathways were identified. MSigDB was used to identify the top 15 enriched biological pathways (padj < 0.05). RT-qPCR was performed to validate changes in mRNA expression of DEG common to both BM-MSC exposures. Given robust upregulation of *CCL2* mRNA expression in HESCs exposed to the BM- and UC-MSC secretomes, transwell migration and invasion assays were performed to determine the effect of recombinant CCL2 on HESC motility. Statistical significance was defined as *p*<0.05.

**Results:** Indirect co-culture of HESCs with BM- or UC-MSCs resulted in significant increase in HESC migration and invasion regardless of the source of MSCs. However, effects on cellular proliferation varied among the MSC donors. Exposure of HESCs to the secretome of BM-MSCs changed the expression of 10,139 genes with FDR < 0.05. There was overlap among 4350 genes between HESCs exposed to BM-MSC-1 and BM-MSC-2. Within four biological pathways enriched in HESCs, 4 genes (*CCL2, HGF, PLAU*, and *BDKRB2*) were differentially expressed in HESCs that had been cocultured with BM-MSC-1 and BM-MSC-2. qRT-PCR showed significantly increased mRNA expression of *CCL2* in HESCs exposed to BM-MSC-1 (5-fold) and BM-MSC-2 (7.7-fold). In contrast, the increase in *HGF* expression was significant after exposure to BM-MSC-2 (1.8-fold) but not BM-MSC-1. Exposure to the UC-MSC secretome had similar effects on HESC-derived *CCL2* and *HGF* levels. *CCL2* expression was significantly increased (6.5-fold) by UC-MSC-2 but not by UC-MSC-1; *HGF* expression was significantly increased (1.6-fold) by UC-MSC-2 but not by UC-MSC-1. Validation studies indicated that exposure to recombinant CCL2 for 48 hours significantly increased HESC migration (1.2-fold) and invasion (1.4-fold). These data suggest that CCL2 is a key factor in mediating MSC-induced HESC motility.

**Conclusion:** Increased HESC motility by the secretome of BM- and UC-MSC appears to be mediated by paracrine and autocrine mechanisms, in part by upregulated *CCL2* expression in HESC. Together, our data support the potential for leveraging the MSC secretome as a novel cell-free therapy in the treatment of disorders of endometrial regeneration.

## Introduction

Cyclic regeneration and renewal of the uterine endometrium is required for human reproductive function. In normal menstrual cycles, demise of the ovarian corpus luteum and the subsequent decrease in circulating progesterone triggers an inflammatory response that results in shedding of the functionalis layer of the endometrium. Regeneration of the functionalis layer after menstruation, and parturition, involves inflammation, angiogenesis, re-epithelization, and tissue remodeling, which are processes integral to wound repair in non-reproductive organs (1). Cyclic repair of the endometrium requires intact function of multiple cell types, including parenchymal, endothelial and immune cells, and contributions from adult stem cells which may originate from endometrial and/or extrauterine sources (2-4).

Mesenchymal stem cells (MSCs) are multipotent adult cells showing multilineage differentiation into cells such as adipocytes, chondrocytes, and osteoblasts. MSCs have been demonstrated to regenerate tissue and to migrate towards injured sites where they can secrete bioactive products to aid in tissue regeneration and repair (5). These secreted factors include soluble factors such as cytokines and insoluble microvesicles (6). The secretome regulates cellular processes critical for tissue regeneration by modulating biological processes such as immunomodulation, angiogenesis, cell migration and proliferation (7). MSCs can be harvested from several anatomical sites including bone marrow (BM), adipose tissue, placenta, and umbilical cord. BM-derived MSCs (BM-MSCs) are the most frequently used stem cells in tissue regeneration and may be considered the “gold standard”, however, MSCs derived from other sources also possess the ability to home to injured sites and facilitate tissue repair (8-11).

Asherman syndrome, characterized by intrauterine adhesions and inadequate endometrial regeneration, is a significant cause of female infertility with limited therapeutic options. Studies with rodent models and small clinical trials have shown that non-hematopoietic MSCs from different tissue sources can contribute to endometrial repair after injury (12). In rat models of endometrial injury, BM-MSCs and umbilical cord (UC) MSCs aid endometrial repair through paracrine actions on the endometrium as well as differentiation into endometrial cell types (13-16). The secretome and biological characteristics of MSCs vary based on tissue source (17, 18); however, both BM-MSCs and UC-MSCs promote endometrial regeneration in rodent models (12). In a small trial of 26 women with intrauterine adhesions, transplantation of UC-MSCs into the uterus after adhesiolysis increased endometrial thickness and expression of factors associated with endometrial proliferation (19). Together, these studies suggest that MSCs can be leveraged as potential primary or adjunct therapeutic approaches to enhance or restore endometrial regeneration (20, 21). However, the efficacy and safety of such cell-based therapies remains unproven, rendering use of the MSC secretome or its components as an attractive potential alternative for cell-free therapy.

Whereas prior studies to determine the effects of MSCs on endometrial regeneration and repair have focused on endometrial cell proliferation, we tested the hypothesis that the MSC secretome enhances endometrial stromal cell motility and regulates expression of genes that promote cell motility. We used a coculture model with human MSCs from BM and UC. This model allowed for the secretome to mediate an interaction with human endometrial stromal cells (HESCs). Our first objective was to determine how exposure to BM-MSCs impacts proliferation, migration, and invasion of HESCs. To identify underlying molecular mechanisms associated with the observed changes in HESCs, we performed RNA sequencing (RNA-Seq) in HESCs exposed to BM-MSCs. Given the noninvasive, easy accessibility, and increasing utilization of human UC-MSCs in regenerative medicine, our second objective was to determine if UC-MSCs and BM-MSCs have a similar effect on HESCs. We found that exposure to the secretome of both UC- and BM-MSCs significantly increases HESC migration and invasion, with a modest effect on HESC proliferation. Increased HESC motility induced by the MSC secretome appears to be mediated by paracrine and autocrine mechanisms, in part by upregulation of HESC CCL2 expression. The potential to leverage the MSC secretome as a novel cell-free therapy in the treatment of disorders of endometrial regeneration is discussed.

## Methods

### Cell lines

hTERT-immortalized human endometrial stromal cells ((HESCs), ATCC) were grown in phenol red-free DMEM/F12 media (Gibco) supplemented with 10% charcoal stripped fetal bovine serum (FBS), 1% ITS+premix (Corning) and 500 ng/mL puromycin (Sigma) and maintained in a humidified incubator under standard culture conditions, as per manufacturer protocols. Human BM-derived mesenchymal stem cells (ATCC) were derived from a 35-year-old Hispanic female donor. These commercially obtained BM-MSCs, designated BM-MSC-1, were cultured in MSC basal media (ATCC) supplemented with MSC Growth kit (ATCC) [hereafter referred to as “MSC media”] as per manufacturer protocols. Cells were used for all experiments prior to passage 6.

### Human subjects

BM aspirates and UC were collected using a protocol approved by the Institutional Review Board of Rutgers Biomedical and Health Sciences. Three women, age 20-30 years, provided written informed consent to undergo BM aspiration for use of their BM cells. Two women, age 20-30 years, provided written informed consent for the procurement and use of placental tissues. All subjects were generally healthy. Specific inclusion criteria for UC samples included cesarean delivery of a full-term male infant. Exclusion criteria for both samples included infectious disease, obese or underweight body mass index, chronic medical conditions, acute medical conditions at time of sample collection, and pregnancy-related complications for UC donors.

### Isolation of BM-MSCs

BM aspirates were collected into heparinized syringes for transport to the laboratory and anonymized as BM-MSC-2, BM-MSC-3, and BM-MSC-4. Unfractionated BM was diluted 1:1 in DMEM with 10% FBS and added to vacuum gas plasma treated plates (BD Falcon). After three days, red blood cells and granulocytes were removed by Ficoll-Hypaque density gradient centrifugation, and the mononuclear cells were returned to the plates with fresh media. At weekly intervals 50% of the media was replaced with fresh media until the adherent cells reached 80% confluence. Adherent cells were serially passaged at 80% confluence until the fourth passage, at which time the cells were confirmed by phenotype and multilineage differentiation potential as described below. Confirmed MSCs were used through passage 6.

### Isolation of UC-MSCs

Human placentas were collected after full-term cesarean delivery of healthy, full-term male infants and transported to the lab in 1x phosphate buffered saline (PBS); placentas were processed within 1 hour or stored at 4°C for up to 18 hours prior to processing. Upon processing the samples were anonymized as UC-MSC-1 and UC-MSC-2. Within 24 hours of delivery the UC was excised from the placenta and washed with PBS to remove blood and mucus, transferred to petri dishes and cut into 1×1 cm pieces. The outermost region of the UC was used for UC-MSC isolation; the Wharton’s jelly, and vein and artery region were discarded. Up to 15-20 UC pieces were transferred to tissue culture dishes, ensuring adequate spacing between each piece, and left without media for 10-15 min to attach. After that time, 10 mL media (Dulbecco’s modified Eagle’s media (DMEM, 1 g/L D-Glucose; Invitrogen) containing 10% fetal bovine serum (FBS; Hyclone) and 1% penicillin/streptomycin (Invitrogen) was added to each culture dish. Medium was fully replaced with fresh medium every 2-3 days until outgrowth from the UC pieces was visible. The sections were then transferred to a new culture dish. Up to 7 serial transfers were performed. Culture dishes with outgrowth were left for ∼1 week until expansion of the cells was observed, this step was counted as passage 0. Adherent cells were serially passaged at 80% confluency and cryopreserved through passage 3. At the fourth passage cells were confirmed to be MSCs by phenotyping and multilineage differentiation capacity as described below. Confirmed MSCs were used through passage 6.

### Characterization of Mesenchymal Stem Cells

Human BM-MSCs and UC-MSCs were phenotyped by flow cytometry. All conjugated antibodies were purchased from Becton Dickinson. MSCs were confirmed to be negative for CD45 and positive for CD44, CD90, CD73, and CD29 expression. All antibodies were used at 1:100 dilution and analyzed using FACScalibur flow cytometer and FlowJo software (Becton Dickinson).

#### Osteogenesis and Von Kossa staining

MSCs were seeded in a 6-well plate at 10,000 cells per cm^2^ in MSC media, and incubated in a 37°C, 5% CO_2_ humidified incubator overnight. The following day the cells were washed with PBS and media changed to human osteoblast differentiation media (Sigma-Aldrich). Osteoblast differentiation media was changed every 3 days for a period of 14 days as per the manufacturer’s guidelines. The mineral deposition of osteoblast cultures was visualized by Von Kossa staining for calcium. Briefly, cells were fixed with 4% PFA in PBS, incubated with 2% silver nitrate solution in the dark, washed with deionized water, and developed with 1% pyrogallol. Cells were counterstained with nuclear fast red (Vector). Osteoblast nodules containing calcium mineral stain black; less developed nodules stain yellow to brown.

#### Adipogenesis and Oil Red O staining

MSCs were seeded in wells of 6-well plates at 40,000 cells/cm^2^ in MSC media and incubated in a 37°C, 5% CO_2_ humidified incubator for 3 days. The cells were washed with PBS and media replaced with human adipocyte differentiation media (Sigma Aldrich). The medium was replaced with fresh human adipocyte differentiation media every 3 days as per the manufacturer’s protocol. Within 3-4 weeks cells were differentiated into adipocytes with large lipid droplets. Cells were fixed with 4% PFA in PBS, washed twice with water and incubated with 60% isopropanol. Cells were then incubated with Oil Red O Working Solution (Sigma Aldrich) and counterstained with hematoxylin prior to visualization with a light microscope.

### Cell proliferation assay

A transwell co-culture system using 0.4µm pore size cell culture inserts was utilized to assess the impact of MSC secretome exposure on HESC proliferation. BM-MSCs, UC-MSCs, and HESCs were cultured in their respective media in 100 mm tissue culture dishes, grown to 70-80% confluency, and then harvested and washed with PBS. HESCs were transferred to 6-well plates containing 12mm diameter glass circles (Fisher) at 2×10⁵ cells per well. BM-MSCs were placed on 6-well plate formatted culture inserts with 0.4µm pore membrane (Corning) at 1×10⁵ cells per insert. BM-MSCs and HESCs were cultured independently for 24 hours in their respective media. All cells were washed with PBS and were then exposed to co-culture media (phenol red-free DMEM supplemented with 10% charcoal stripped FBS and 1% ITS+premix) in the transwell system for 24 hours. HESCs cultured in the transwell system without MSCs served as controls. After 24h, HESCs were fixed with 4% PFA, washed with PBS, permeabilized with PBST (1x PBS with 0.1% Triton-100) and incubated with anti-Ki67 antibody (Abcam, 1:500) for 2 hours at room temperature. Cells were incubated with a secondary antibody, Alexa Fluor 488 (Thermofisher, 1:1000 dilution) for 30 minutes at 4°C. Vectashield with DAPI (Vector) was used for mounting and nuclear visualization. Slides were imaged using the Leica DM2500 LED microscope and Leica Application Suite X (LAS X) software (Leica Microsystems). Two independent observers counted the number of Ki67 positive cells and the total number of cells per field in three 40x magnification fields per slide using ImageJ software. Percentage of proliferating cells was determined as the number of Ki67 positive cells per total cell count × 100.

### MSC and HESC co-culture transwell migration assay

For transwell migration assays, 24-well transwell chambers (Corning) and Falcon transparent cell culture inserts (8 μm pore size, Corning) were used. The upper compartment contained 5×10⁴ HESCs in phenol red-free DMEM supplemented with 0.5% charcoal stripped FBS and 1% ITS+premix. The lower compartment of the chamber contained BM-MSCs or UC-MSCs in phenol red-free DMEM supplemented with 10% charcoal stripped FBS and 1% ITS+premix. HESCs exposed to DMEM supplemented with 10% charcoal stripped FBS and 1% ITS+premix served as the control. The extent of HESC migration was assessed at 24 hours. Non-migrated cells in the upper compartments were removed with a cotton swab. Cells on the undersurface of the membrane were fixed with 4% PFA and stained with H&E. Slides were imaged using the Leica DM2500 LED microscope and Leica Application Suite X (LAS X) software at 2.5x magnification. Two independent observers counted the nuclei. The extent of cell migration was determined by counting nuclei of cells that migrated in a 1250 × 1250 pixel area of the insert using ImageJ software.

### MSC and HESC co-culture transwell invasion assay

Invasion assays were performed in 24-well transwell chambers using 8 μm pore size cell culture inserts (Corning). Cell culture inserts were coated with DMEM media with 0.3 mg/mL Matrigel (phenol red-free, growth factor reduced basement membrane matrix, Corning). Experiments were performed in the exact same manner as described for the transwell migration assays except 1× 10^5^ HESCs were added to the upper compartment of the chambers.

### Next-generation mRNA sequencing

For mRNA-seq experiments, the transwell co-culture system utilized to assess HESC proliferation was used. 2 × 10^5^ HESC cells per well were plated in 6-well dishes (Sigma-Aldrich). As performed for the proliferation assays, BM-MSCs were plated onto 0.4 µm pore size cell culture inserts.

Total RNA was isolated from HESCs with the RNeasy Plus Mini Kit (Qiagen) after 24 hours of co-culture with BM-MSCs and RNA integrity number (RIN) determined. mRNA-Seq library construction was done using NEBNext UltraRNA Library Prep Kit for Illumina (NEB) at Novogene Corporation Inc. The multiplexed library was sequenced on an Illumina Novaseq 6000 platform (Illumina) in a 2 × 150 bp configuration to a target depth of 20–30 M reads per sample.

Sequence analysis was performed following NF-Core RNA-Seq guidelines (v1.4.2) for rigor and reproducibility (22). The raw reads were aligned to the human genome (GRCh37) using STAR (v2.6.1), followed by transcript abundance and hit counts generation through StringTie (v2.0) and featureCounts (v1.6.4) (23, 24). Subsequent hit count normalization and pairwise differential gene expression comparisons were performed using the DESeq2 (v1.22.1) method with significance thresholds set at FDR adjusted *p*< 0.05 (25). Pathway term enrichment was performed via Ingenuity Pathway Analysis. mRNA-Seq data have been uploaded to the Genome Expression Omnibus/National Center for Biotechnology Information with accession number GSE217506.

### Quantitative PCR

We applied the transwell system described for the proliferation studies for RT-qPCR. HESCs at 2 × 10^5^ per well were plated in 6-well dishes (Sigma-Aldrich), with BM-MSCs or UC-MSCs in the inner wells with 0.4 µm pore size inserts. After 24 h, total RNA was isolated from HESCs using the RNeasy Plus Mini Kit (Qiagen). Total RNA (1 µg) was reverse transcribed using qScript cDNA Supermix (Quanta BioSciences). RT-qPCR was performed in triplicate using the QuantiNova SYBR Green PCR Kit (Qiagen) and the following primer sequences: *18s* forward 5’-CCG GGC TTC TAT TTT GTT GGT-3’, *18s* reverse 5’-TAG CGG CGC AAT ACG AAT G-3’, *CCL2* forward 5’-GCT CAT AGC AGC CAC CTT CAT TC-3’, *CCL2* reverse 5’ GCT CAT AGC AGC CAC CTT CAT TC-3’, HGF-1 forward 5’-GCA ATT AAA ACA TGC GCT GAC A-3’, HGF-1 reverse 5’-TCC CAA CGC TGA CAT GGA AT-3’. The relative expression level of each target gene was quantified using (2^−ΔΔCt^) comparing to 18s rRNA (26). Expression level was determined as a mean of the triplicates.

### Recombinant (r) CCL2 on HESC migration and invasion

To assess the effect of rCCL2 on HESC motility, transwell migration and invasion assays were performed using 24-well transwell chambers (Corning) and Falcon transparent cell culture inserts (8 μm pore size, Corning). To assess HESC migration, 5×10⁴ HESCs in phenol red-free DMEM/F12 media supplemented with 0.5% charcoal stripped FBS, 1% ITS+premix and 100 ng/ml recombinant CCL2 protein (R&D Systems) or 1% bovine serum albumin (BSA) for the control were placed on uncoated inserts in the upper compartment. To assess HESC invasion, 1×10^5^ HESCs in phenol red-free DMEM/F12 media supplemented with 0.5% charcoal stripped FBS, 1% ITS+premix and 100ng/ml recombinant CCL2 protein or 1% BSA for the control were placed on Matrigel coated inserts in the upper compartment. Phenol red-free DMEM/F12 media supplemented with 10% charcoal stripped FBS and 1% ITS+premix was added to the lower compartment of the chamber. After 24 hours, non-migrated/non-invaded cells in the upper compartments were removed with a cotton swab. Cells on the undersurface of the membrane were fixed with 4% PFA and stained with H&E. Slides were imaged using the Leica DM2500 LED microscope and Leica Application Suite X (LAS X) software at 2.5x magnification. Two independent observers counted the nuclei. The extent of cell migration/invasion was determined by counting nuclei of cells that migrated in a 1250 × 1250 pixel area of the insert using ImageJ software.

To assess HESC migration and invasion after 48 hours, 2×10⁴ HESCs in phenol red-free DMEM/F12 media supplemented with 1% charcoal stripped FBS, 1% ITS+premix and 100ng/ml recombinant CCL2 protein (R&D Systems) or 1% BSA for the control were placed on uncoated or Matrigel coated inserts in the upper compartment. Phenol red-free DMEM/F12 media supplemented with 5% charcoal stripped FBS and 1% ITS+premix was added to the lower compartment of the chamber. After 48 hours, non-migrated/non-invaded cells in the upper compartments were removed with a cotton swab. Cells on the undersurface of the membrane were fixed with 4% PFA and stained with H&E. Slides were imaged using the Leica DM2500 LED microscope and Leica Application Suite X (LAS X) software at 2.5x magnification. Two independent observers counted the nuclei. The extent of cell migration/invasion was determined by counting nuclei of cells that migrated in a 1250 × 1250 pixel area of the insert using ImageJ software.

### Statistical analysis

All experiments, with exception of mRNA-Seq, were performed at least three times with at least three technical replicates. Means were compared using unpaired t-test or ANOVA followed by Tukey’s multiple comparison test. Data are expressed as mean ± SEM. Statistical analyses were performed using Prism Version 8.0 (GraphPad). Statistical significance was defined as *P* < 0.05.

## Results

### Characterization of BM-MSCs

MSCs are mesoderm-derived, multipotent adult stem cells that can differentiate into a variety of cell types. We characterized BM-MSCs by phenotype using established surface markers (27). Flow cytometry indicated dim fluorescence on <1% of cells expressing the hematopoietic marker CD45 (Fig. 1A and B). For all BM-MSCs (1-4), >80% expressed CD44, CD73, CD29 and CD90 (Fig. 1A and B). (Data are not shown for BM-MSC-3 and BM-MSC-4.) Next, we performed multilineage differentiation assays and showed adipogenic differentiation (Fig. 1C and D) as well as osteogeneic differentiation (Fig. 1F and G).

**Figure 1.**
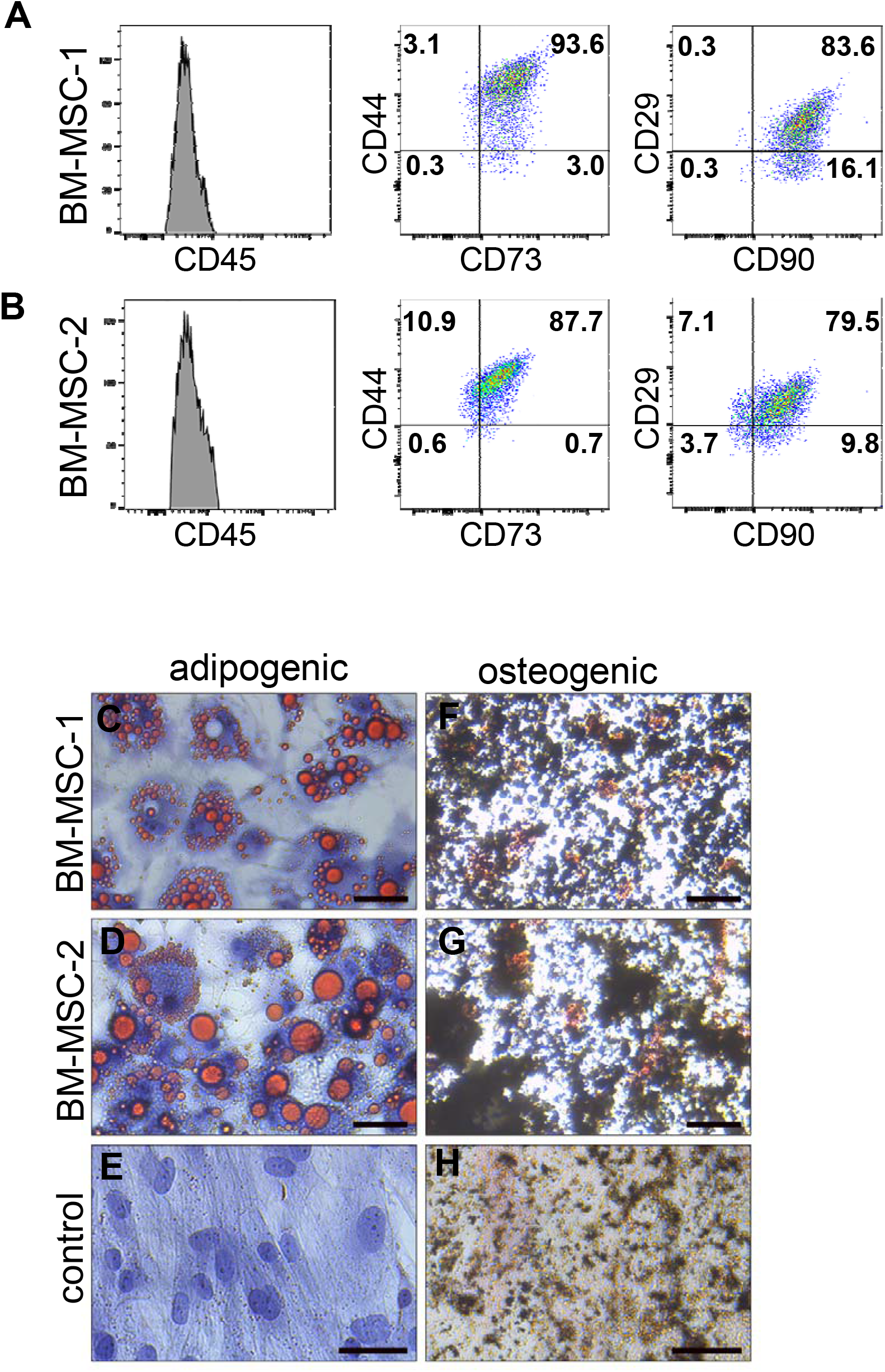
Surface marker expression and multipotent differentiation of BM-MSCs. (**A, B**) Flow cytometric analysis was performed on (A) BM-MSC-1 and (B) BM-MSC-2. MSC surface markers CD44, CD73, CD29, and CD90 were highly expressed, whereas expression of hematopoietic stem cell marker CD45 was not detected. The percentages of cells expressing each surface marker are indicated. (**C, D**) Following culture of BM-MSCs in adipocyte differentiation media for 21 days, Oil Red O staining was used to detect adipocytes. Lipid droplets appear red, and nuclei appear blue. (**E**) BM-MSC-1 cultured in control MSC media for 21d and stained with Oil Red O. (**F, G)** Following exposure of BM-MSCs to osteoblast differentiation media for 30 days, von Kossa staining was used to identify osteoblasts. Bone nodules containing calcium mineral stain black. (**H**) BM-MSC-1 cultured in control MSC media for 30d and stained with von Kossa stain. Scale bars: 50 mm.

### The BM-MSC secretome mediated HESC migration and invasion

We used transwell coculture systems to assess the role of the BM-MSC secretome on HESC motility (Fig. 2A). To determine if exposure to the BM-MSC secretome enhances HESC migration toward a chemoattractant, transwell migration of HESCs across an 8 µm pore transwell chamber culture insert was determined. BM-MSCs were cultured in media supplemented with 10% FBS in the lower compartment. HESCs resuspended in media supplemented with 0.5% FBS were placed in the upper compartment of the transwell chamber, and after 24 h, the number of migrated HESCs was counted (Fig. 2A and B). Indirect coculture with BM-MSCs significantly increased HESC transwell migration, 3.9-fold for BM-MSC-1, 7.3-fold for BM-MSC-2, 5.8-fold for BM-MSC-3 and 3.4-fold for BM-MSC-4, relative to culture in media without BM-MSCs (Fig. 2C and Supplementary Fig.1A).

**Figure 2.**
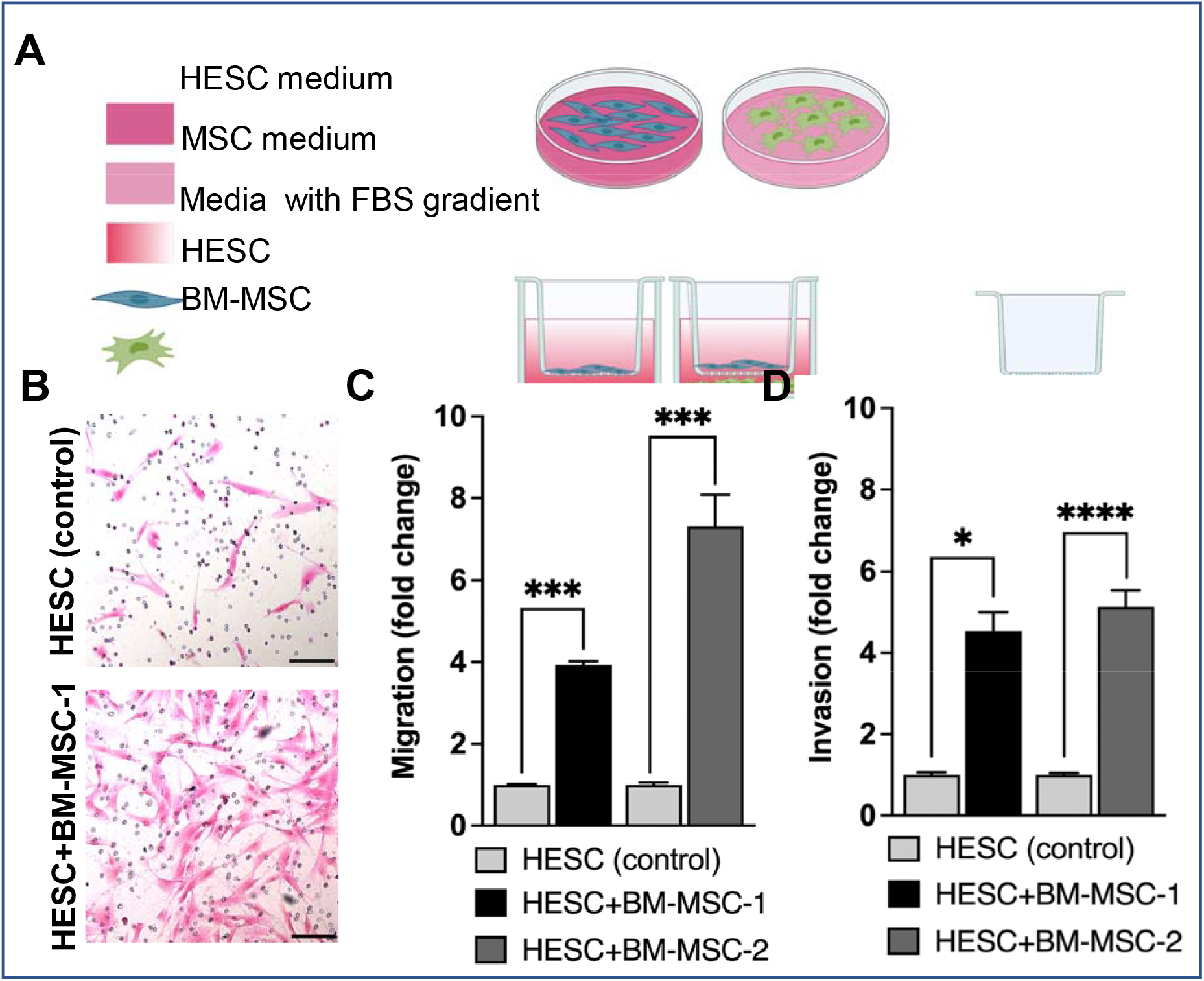
The BM-MSC secretome increases HESC migration and invasion. HESC transwell migration and invasion through Matrigel were assessed after indirect coculture with BM-MSCs for 24h. (**A**) Diagram of the coculture system of HESCs with BM-MSCs used to assess HESC migration and invasion. (**B**) Representative images of H&E-stained HESCs that migrated or invaded after 24h coculture. Scale bars 100µM. (**C**) Transwell migration was significantly increased by coculture with BM-MSC-1 (\*\*\**P*=0.0009) and BM-MSC-2 (\*\*\**P*=0.0004) compared to control HESCs. (**D**) Transwell invasion through Matrigel was significantly increased by coculture with BM-MSC-1 (\**P* =0.02) and BM-MSC-2 (\*\*\*\**P*< 0.0001) compared to control HESC. Data are expressed as means (of N=3–6) ± SEM. Statistical analysis was performed with an unpaired t test followed by Welch’s correction.

We next asked if the secretome of BM-MSCs could enhance HESC invasion through a Matrigel matrix. As in the transwell migration assays, a 20-fold increase in FBS was used as a chemoattractant. 24h of indirect coculture with BM-MSC-1, BM-MSC-2, and BM-MSC-3 significantly increased HESC transwell invasion, 4.5-fold, 5.1-fold, and 4.2-fold, respectively relative to culture in media without BM-MSCs (Fig. 2D and Supplementary Fig.1B). HESC invasion after coculture with BM-MSC-4 was 2.8-fold greater than control, but this increase was not statistically significant (Supplementary Fig.1B). Together, these data show that BM-MSCs from all donors can enhance HESC motility.

### BM-MSC secretome increases HESC proliferation

We used a transwell coculture system to determine if exposure of HESC to BM-MSC secretome increased HESC proliferation (Fig 3A). HESCs were placed in the lower compartment of the transwell chamber, and BM-MSCs in the inner chambers with 0.4 µm insert. After 24 h, the HESCs were assessed for proliferation based on the percentage of cells expressing Ki67 (Fig. 3B and C). The results indicated that BM-MSC-1 secretome showed a larger percentage of HESCs expressing Ki67 as compared to HECSs cultured in control media (53.8 ± 3.3% vs. 42.7 ± 2.8%, *P*=0.02) (Fig. 3C). These findings indicated that exposure to the BM-MSC-1 secretome increased HESC proliferation. In contrast, we noted no significant (*P*>0.05) difference in the percentage of HESCs expressing Ki67 after exposure to BM-MSC-2 vs. control media (41.6 ± 3.6% vs. 46.1 ± 3.5%) (Fig. 3C). Together, these data show that effect of the BM-MSC secretome on HESC proliferation might depend on the MSC donor.

**Figure 3.**
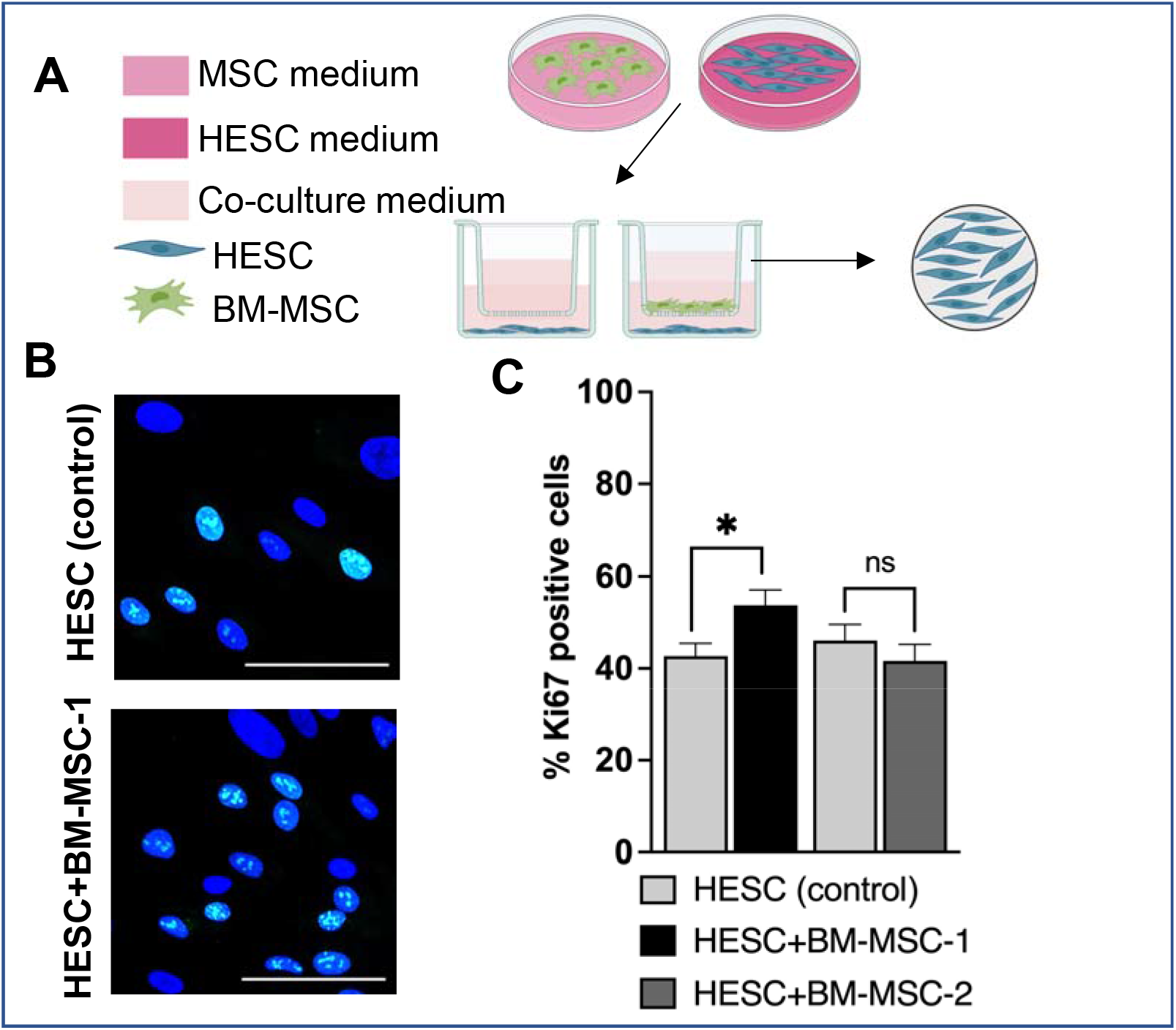
Indirect coculture of HESC with BM-MSC affects HESC proliferation. HESCs were analyzed for cellular proliferation using Ki67 after 24h indirect coculture with BM-MSCs. (**A**) Diagram of the coculture system of HESCs with BM-MSCs used to assess HESC proliferation. (**B)** Representative IF images of HESCs stained with Ki67. Scale bars: 100µM. (**C**) Coculture with BM-MSC-1 significantly increased HESC proliferation (\**P*=0.02). HESC proliferation was unchanged with exposure to BM-MSC-2.

### BM-MSC secretome upregulates cell motility genes

To examine the impact of the BM-MSC secretome on HESC gene expression, whole-genome transcriptome profiling (mRNA-seq) was performed in HESCs cocultured with BM-MSCs in a transwell system. We sought to identify changes in HESC gene expression after 24 h of transwell coculture with BM-MSC-1 (commercially purchased) or BM-MSC-2 (MSCs isolated in our laboratory) relative to HESCs cultured in control media. PCA dimensionality reduction of the mRNA-seq data showed that the HESCs clustered into three groups (Fig. 4A). mRNA-seq analysis revealed 10,139 differentially expressed genes (DEGs) with FDR < 0.05 (Fig. 4B). Of these, 4160 DEGs were identified in HESC exposed to BM-MSC-1, 1629 DEGs in HESC exposed to BM-MSC-2, and 4350 DEGs overlapped in HESC exposed to BM-MSC-1 and BM-MSC-2 (Fig. 4B). DEGs with *Padj* <0.05 and FC > 1.5 or FC < -1.5 were analyzed with Ingenuity Pathway Analysis (IPA) to identify enriched pathways with -log (p-value) > 1.3. After exposure to the BM-MSC secretomes, 16 pathways were enriched in HESC+BM-MSC-1 and 78 pathways were enriched in HESC+BM-MSC-2 (Supplementary Tables 1 and 2). Four pathways, *Coagulation system, NAD Signaling Pathway, Renal Cell Carcinoma Signaling* and *Tumor Microenvironment Pathway* were enriched in HESCs exposed to the secretomes from both BM-MSC-1 and BM-MSC-2 (Fig. 4D). Within those 4 pathways, only 4 genes (*CCL2, HGF, PLAU*, and *BDKRB2*) were differentially expressed in HESCs that had been cocultured with both sources of BM-MSCs (Fig. 4E). We used qRT-PCR to confirm the increased expression of *CCL2* and *HGF* in HESCs exposed the BM-MSC secretome. qRT-PCR showed significantly increased mRNA expression of *CCL2* in both HESC+BM-MSC-1 (5-fold) and HESC+BM-MSC-2 (7.7-fold) compared to control HESC (Fig. 4F). In contrast, the increase in *HGF* expression was significant in HESC+BM-MSC-2 (1.8-fold), but not significantly different in HESC+BM-MSC-1 compared to control HESCs (Fig. 4G).

**Figure 4.**
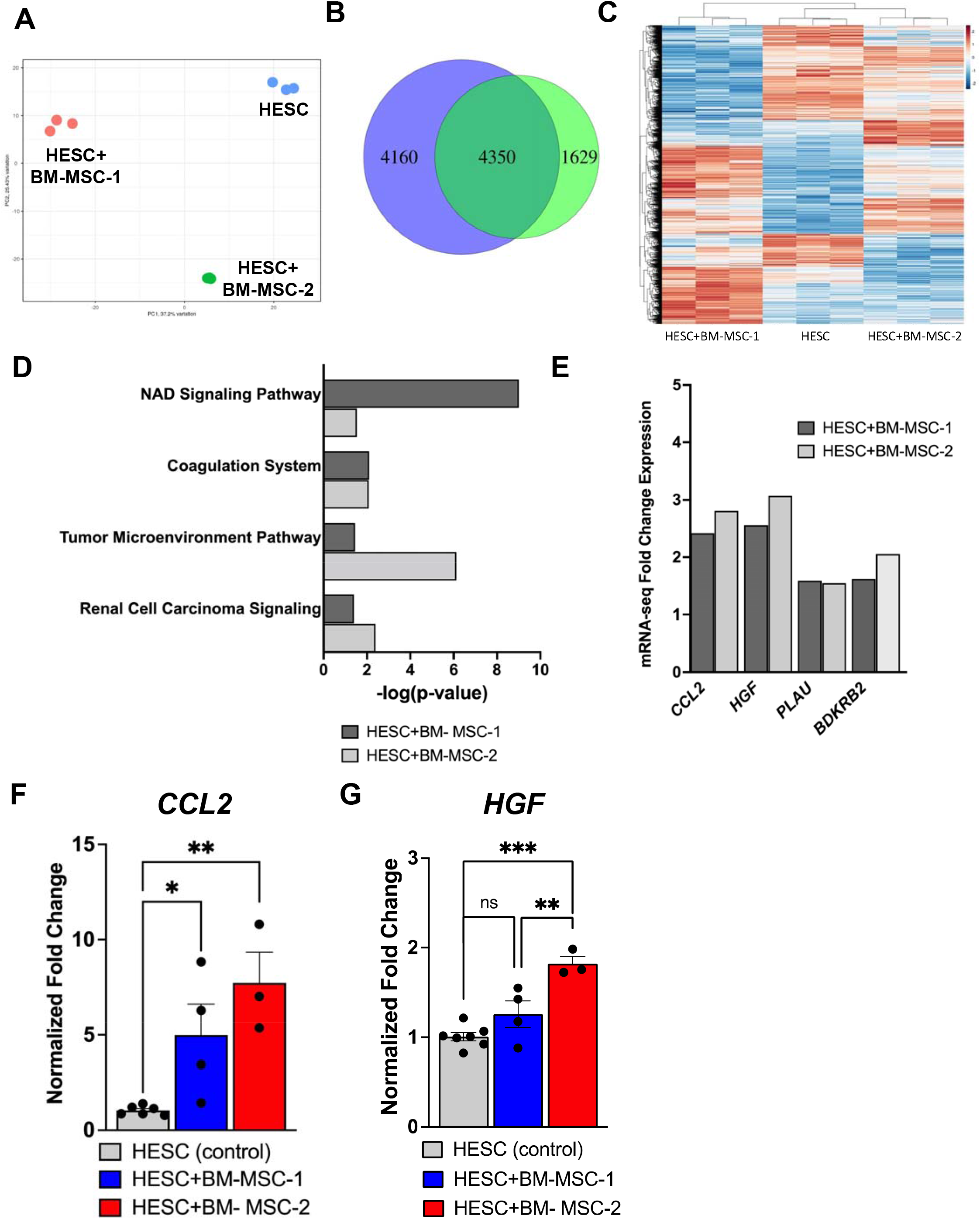
Gene expression changes in HESCs after coculture with BM-MSC-1 and BM-MSC-2. (**A**) Principal component analysis results of the transcriptome from mRNA-seq data of control HESCs and HESCs cocultured with BM-MSCs (HESC+BM-MSC-1 and HESC+BM-MSC-2). (**B**) Venn diagram illustrating number of differentially expressed genes (DEGs) identified in HESCs exposed to BM-MSC-1 (purple), BM-MSC-2 (green), and number of DEGs common to both BM-MSC exposures (blue overlap). **(C)** The heat map of all DEGs in control HESCs and HESCs cocultured with BM-MSCs. The X-axis represents the different samples (N=3, each condition) and the Y-axis shows all 10,139 DEGs. The color key from orange to blue indicates the relative gene expression level from high to low. (**D**) Ingenuity Pathway Analysis (IPA) identified four significant canonical signaling pathways with -log(p-value) > 1.3 in HESCs cocultured with BM-MSC-1 and BM-MSC-2. (**E**) Four DEGs within the four canonical pathways identified in HESCs cocultured with BM-MSC-1 and BM-MSC-2. (**F**,**G**) RT-qPCR was used to determine expression of *CCL2* and *HGF* in HESCs cocultured with BM-MSCs. The relative expression was compared to 18s rRNA. (**F**) Expression of *CCL2* was significantly increased in HESC+BM-MSC-1 (N=4, \**P*=0.04) and HESC+BM-MSC-2 (N=3, \*\**P*=0.004) compared to control HESC (N=6). (**G**) Expression of *HGF* was unchanged in HESC+BM-MSC-1 (N=4) and significantly increased in HESC+BM-MSC-2 (N=3, \*\*\**P*=0.0002) compared to control HESC (N=6). Expression of *HGF* was significantly increased in HESC+BM-MSC-2 compared to HESC+BM-MSC-1 (\*\**P*= 0.006). Data are expressed as means ± SEM. Statistical analysis was performed with ANOVA followed by Tukey’s multiple comparisons test.

Together, these analyses reveal DEGs and pathways that are similarly altered in HESCs exposed to BM-MSCs from different donors, with *CCL2* as the gene that is consistently significantly upregulated.

### Characterization of UC-MSCs

UC-MSCs are more readily accessible for therapeutic application and have been tested in the uterus (19). We were interested in determining if the secretome of UC-MSCs had the same effects on HESC proliferation, migration, and invasion as the secretome of BM-MSCs. We isolated UC-MSCs from two donors, UC-MSC-1 and UC-MSC-2. First, we used flow cytometry to characterize surface expression of MSC markers. Fewer than 1% of the cells expressed CD45, and greater than 99% of the cells in both UC-MSC-1 and UC-MSC-2 expressed CD44, CD73, CD29 and CD90 (Fig. 5A and B). Next, we performed multilineage differentiation assays to demonstrate multipotency of the UC-MSCs. After 21-28 days of exposure of UC-MSCs to adipocyte differentiation media, differentiated cells contained red stained lipid vacuoles (Fig. 5C and D). After 30 days of exposure of UC-MSCs to osteoblast differentiation media, osteoblasts were identified by mineral deposition that was visualized by Kossa staining for calcium (Fig. 5F and G).

**Figure 5.**
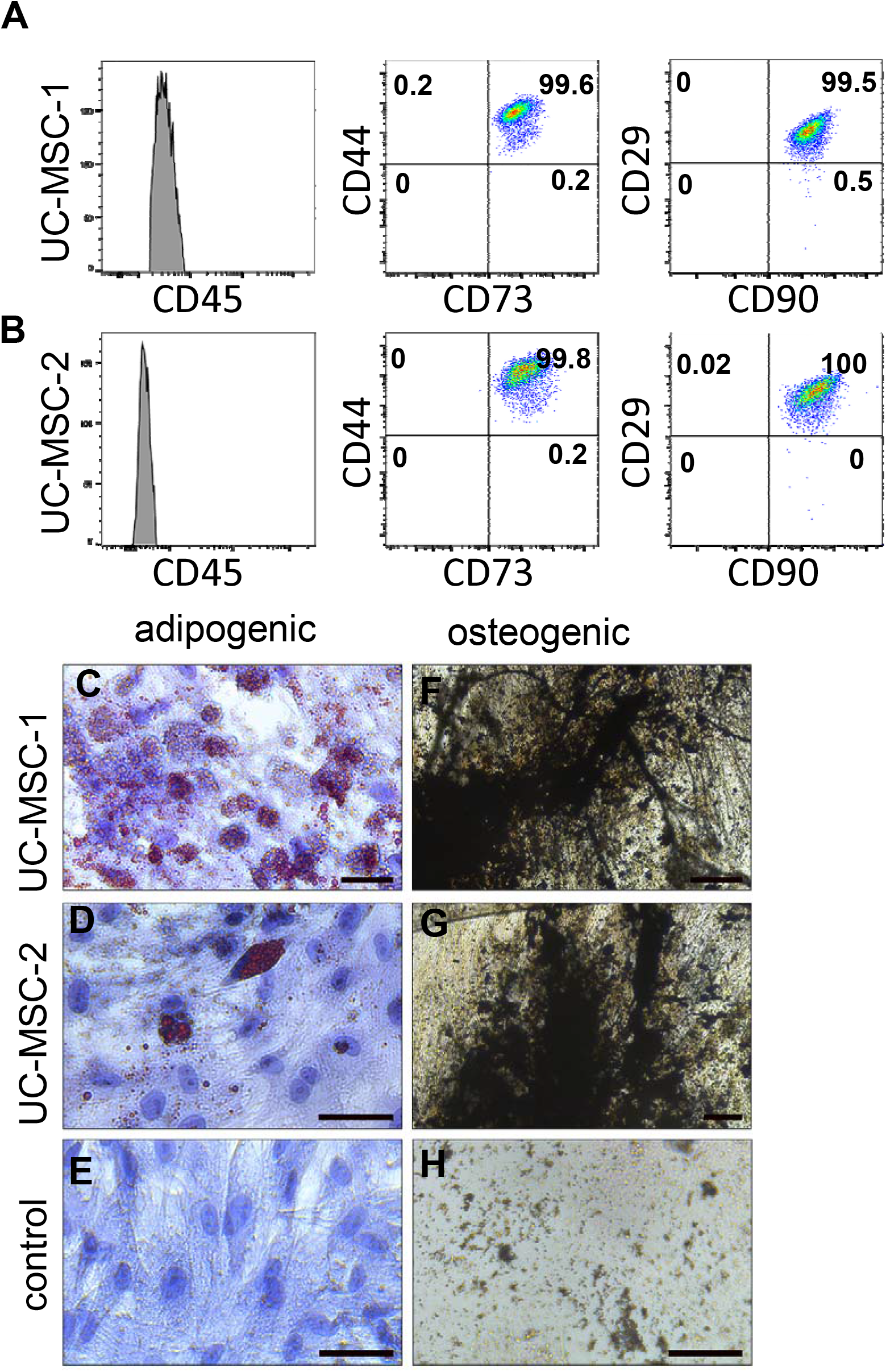
Surface marker expression and multipotent differentiation of UC-MSCs. (**A, B**) Flow cytometric analysis was performed on UC-MSC-1 and UC-MSC-2. MSC surface markers CD44, CD73, CD29, and CD90 were highly expressed, whereas expression of hematopoietic stem cell marker CD45 was not detected. The percentages of cells expressing each surface marker are indicated. (**C, D**) Following culture of UC-MSCs in adipocyte differentiation media for 21 days, Oil Red O staining was used to detect adipocytes. Lipid droplets appear red, and nuclei appear blue. (**E**) UC-MSC-1 cultured in control MSC media for 21d and stained with Oil Red O. (**F, G)** Following exposure of UC-MSCs to osteoblast differentiation media for 30 days, von Kossa staining was used to identify osteoblasts. Bone nodules containing calcium mineral stain black. (**H**) U-MSC-1 cultured in control MSC media for 30d and stained with von Kossa stain. Scale bars: 50 mm.

### Exposure to UC-MSC increases HESC proliferation, migration and invasion

We used transwell coculture systems to determine the impact of the UC-MSC secretome on HESC proliferation (Fig. 3A). We found that the percentage of HESCs expressing Ki67 was similar after 24h coculture with UC-MSC-1 or culture in DMEM+10% charcoal stripped FBS (48.4 ± 1.8% vs. 43.0 ± 3.2%) (Fig. 6A). In contrast, a larger percentage of HESCs expressed Ki67 after coculture with UC-MSC-2 (53.6 ± 1.9% vs. 43.0 ± 3.2%, *p*=0.01) (Fig. 6A). We used the transwell coculture systems with FBS as a chemoattractant to determine if the UC-MSC secretome could enhance growth factor induced HESC migration and invasion (Fig. 2A). We found that coculture with UC-MSCs significantly increased HESC transwell migration and invasion through Matrigel matrix, 3.8-fold for UC-MSC-1 and 3.4-fold for UC-MSC-2 for migration (Fig. 6B) and 3.0-fold for UC-MSC-1 and 4.3-fold for UC-MSC-2 for invasion (Fig. 6C). Thus, like our findings with BM-MSCs, UC-MSC secretome from both donors significantly increased HESC migration and invasion, whereas the secretome from only one donor led to a small, but significant increase in HESC proliferation.

**Figure 6.**
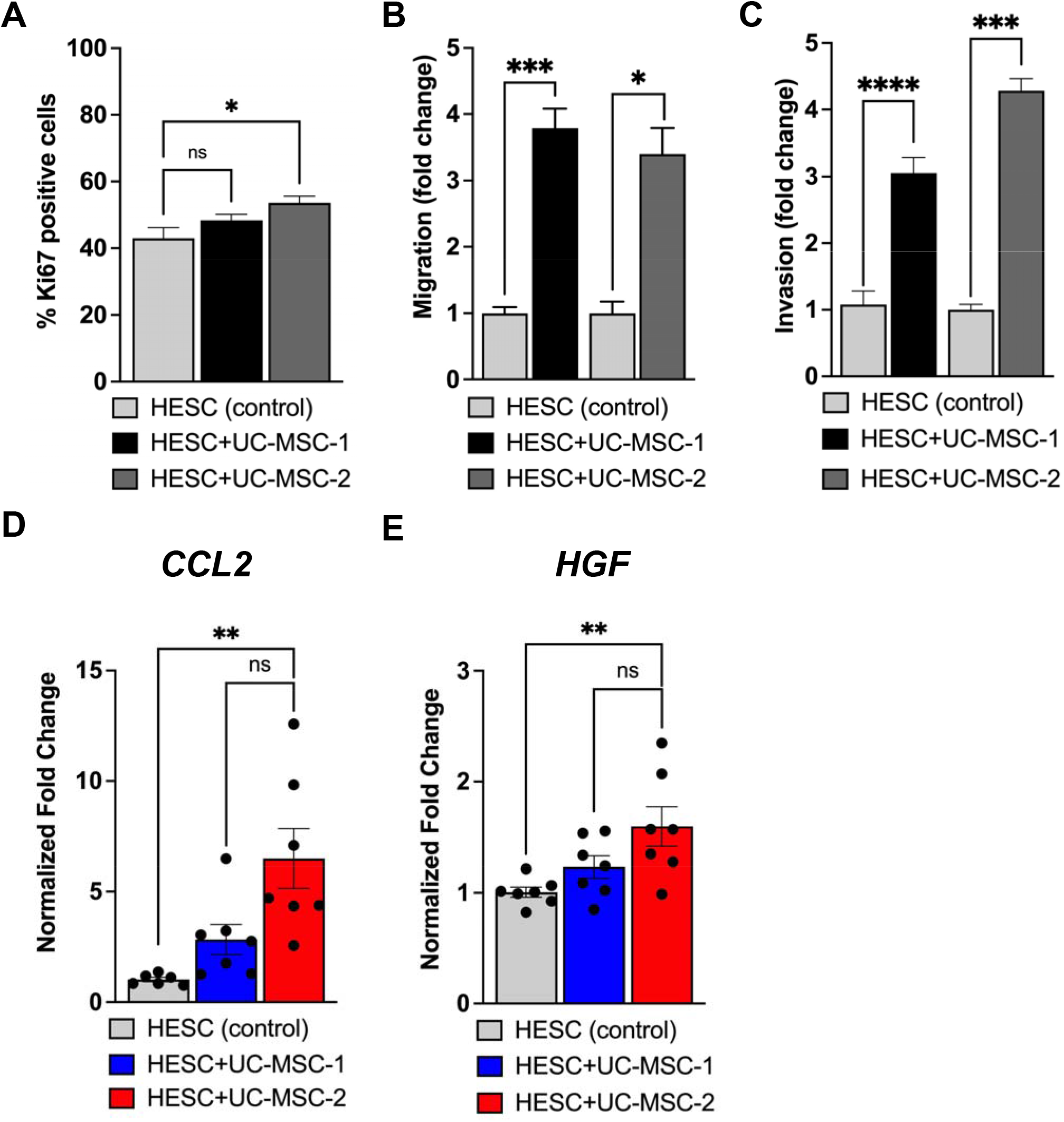
Indirect coculture of HESCs with UC-MSCs affects HESC proliferation, motility and gene expression. **(A)** After coculture with UC-MSCs for 24h, HESCs were analyzed for cellular proliferation using Ki67. HESC proliferation was unchanged with exposure UC-MSC-1. Coculture with UC-MSC-2 significantly increased HESC proliferation (\**P*=0.01). (**B**) HESC transwell migration was assessed after coculture with UC-MSCs for 24h. Transwell migration was significantly increased by coculture with UC-MSC-1 (N=5, \*\*\**P*=0.0004) and UC-MSC-2 (N=3, \**P*=0.01) compared to control HESC (N=5). (**C**) Transwell invasion through Matrigel was significantly increased by coculture with UC-MSC-1 (N=6, \*\*\*\**P*< 0.0001) and UC-MSC-2 (N=3, \*\*\**P*=0.0008) compared to control HESC (N=6) (**D, E**) RT-qPCR was used to determine expression of *CCL2* and *HGF* in HESCs cocultured with UC-MSCs. The relative expression was compared to 18s rRNA. (**D**) Expression of *CCL2* was unchanged in HESC+UC-MSC-1 (N=7) and significantly increased in HESC+UC-MSC-2 (N=7, \*\**P*=0.005) compared to control HESC (N=6). (**E**) Expression of *HGF* was unchanged in HESC+BM-MSC-1 (N=7) and significantly increased in HESC+UC-MSC-2 (N=7), \*\**P*=0.008) compared to control HESC (N=6). Data are expressed as means (of N=3–6) ± SEM. Statistical analysis was performed with ANOVA followed by Tukey’s multiple comparisons test (**A, D, E**) or an unpaired t test followed by Welch’s correction (**B, C**).

### Expression of *CCL2* and *HGF* in HESCs exposed to the UC-MSC secretome

Given that BM-MSCs and UC-MSCs had a similar effect on HESC motility we sought to determine if gene expression changes were similar. We performed RT-qPCR to assess whether indirect coculture with UC-MSCs increased expression of *CCL2* and *HGF*. Expression of *CCL2* in HESCs was significantly increased (6.5-fold) by UC-MSC-2 but not by UC-MSC-1 (Fig. 6D). Expression of *HGF* in HESCs was significantly increased (1.6-fold) by UC-MSC-2, but not by UC-MSC-1 (Fig. 6E). Thus, UC-MSC secretome-induced and BM-MSC secretome-induced changes in *CCL2* and *HGF* expression in HESCs were similar. The robust induction in *CCL2* gene expression, consistently by secretome of BM-MSC from both BM donors tested, and by UC-MSC secretome (1 of 2 donors) suggests that induction of CCL2 is a key component conserved in paracrine signaling from BM-MSCs and UC-MSCs.

### Recombinant CCL2 enhances HESC motility

The BM-MSC secretome is a rich source of many factors, including CCL2 (28). As an autocrine factor, CCL2 can regulate the biological behaviors of endometrial stromal cells in an autocrine manner (29). We hypothesized that CCL2 in the MSC secretome increases secretion of CCL2 by HESCs, resulting in increased HESC motility. To address this hypothesis, we cultured HESCs in media with and without recombinant CCL2 and determined the extent of HESC transwell migration and invasion through Matrigel towards a chemoattractant (Fig 7). We found that exposure to CCL2 for 24 hours, with a 20-fold increase in FBS in the lower compartment as a chemoattractant, did not impact HESC transwell migration or invasion through Matrigel matrix (Fig. 7A, B). We then sought to determine if extended culture with CCL2 could impact HESC motility. For the extended 48 culture, we reduced the initial number of HESCs from 5×10^4^ to 2×10^4^ and used a 5-fold increase in FBS in the lower compartment as a chemoattractant. We found that exposure to CCL2 for 48 hours significantly increased HESC migration (1.2-fold) and invasion through Matrigel (1.4-fold) (Fig. 7A, B). Together, our data suggest that CCL2 is a key factor in mediating MSC-induced HESC motility.

**Figure 7.**
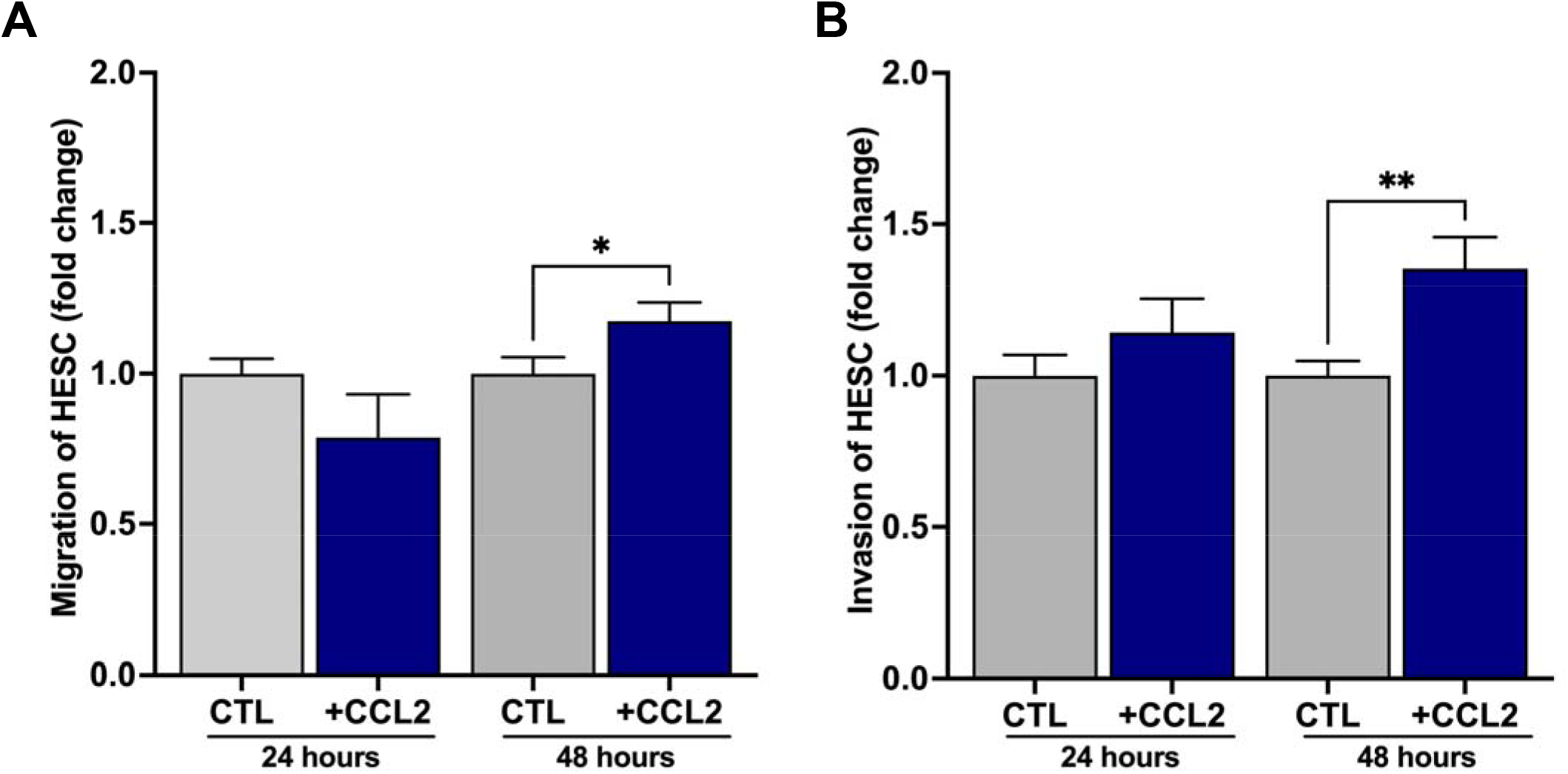
Recombinant CCL2 enhances HESC motility. HESC transwell migration and invasion through Matrigel were assessed after addition of recombinant CCL2 to the culture media. (**A**) At 24 hours, addition of CCL2 did not affect HESC migration. At 48 hours, HESC transwell migration was significantly higher for HESCs exposed to CCL2 compared to control (\**P*=0.05). (B) At 24 hours, addition of CCL2 did not affect HESC invasion through Matrigel. At 48 hours, HESC transwell invasion was significantly higher for HESCs exposed to CCL2 compared to control (\*\**P*=0.005). Data are expressed as means (of N=3–6) ± SEM. Statistical analysis was performed with an unpaired t test.

## Discussion

The normal human endometrium is a highly dynamic tissue that undergoes massive tissue remodeling on a cyclic basis, characterized by repeated tissue breakdown followed by rapid post-menstrual regeneration. In this sense, the endometrium can be considered a “wound” at menses which undergoes cyclic repair, unique in its ability to do so without formation of a scar (30), but the cellular and molecular mechanisms underlying cyclic “scarless endometrial repair” are not completely known. Cyclic repair of the endometrium requires intact function of its multiple cell types, including parenchymal, endothelial and immune cells, as well as contributions from adult stem cell types, including mesenchymal stem cells which may originate from endometrial and/or extrauterine sources (2-4).

In many tissues, mesenchymal stem cells, including those derived from bone marrow, induce rapid repair of damaged tissue (11, 31, 32). Given the central role of endometrial stromal cell motility in the substantial tissue remodeling associated with endometrial regeneration (33), and the well-established role of MSCs in tissue regeneration/repair, we investigated the impact of the MSC secretome on endometrial stromal cell proliferation, motility, and invasive capacity, functions necessary for cyclic endometrial regeneration and repair. MSCs are non-hematopoietic, multipotent adult stem cells which traffic to sites of injury to repair damage by interacting with the niche and releasing secretome such as soluble factors (e.g. chemokines, cytokines, other growth factors) and small microvesicles such as exosomes (5, 31). The MSC secretome mediates tissue repair in a variety of diseased and injured tissues (e.g. cardiac, liver) by mediating angiogenesis and fibrosis pathways (34-36). We focused on effects of MSC derived from bone marrow (BM-MSC) as well as umbilical cord (UC-MSC) given relative ease in accessibility, distinct secretome profiles, and well-established roles for their respective secretomes in tissue remodeling and repair (6). We found that exposure of HESCs to the secretome of both MSC types, from multiple donors, consistently and significantly increased HESC migration and invasive capacity. In contrast, effect of the UC- and BM-MSC secretome on increasing HESC proliferation was modest and donor-dependent. The BM-MSC secretome enriched HESC expression of genes involved in multiple cell signaling pathways involved in cell motility, with a robust and significant increase in CCL2 expression in HESCs exposed to MSCs from UC and BM. This finding, coupled with significant upregulation of HESC motility in response to recombinant CCL2, indicates the upregulation of HESC CCL2 expression as one mechanism by which the MSC secretome enhances HESC motility.

Our findings have significant implications regarding mechanisms by which MSCs may support endometrial regeneration in both physiologic and pathologic settings. Although highly efficient in its physiologic repair processes after menses and parturition, a significant injury may result in intrauterine adhesions and/or atrophy due to inadequate repair. Severe intrauterine adhesions (Asherman syndrome) and endometrial atrophy contribute to female infertility and are among the most difficult causes of infertility to treat, given a lack of therapeutic options. BM-MSCs traffic to the endometrium (37) and have been implicated in endometrial repair in rodent models of Asherman syndrome (38, 39), but the precise mechanisms remain unknown. Although a number of studies provide evidence supporting the ability of BM-derived stem cells to differentiate into endometrial cell types in both rodent (40-42) and human models (41, 43), the overall number of endometrial cells of BM origin is very small. On the contrary Ong et al. (44) found no evidence of transdifferentiation in a chimeric mouse model; rather, macrophages of BM-origin which weakly expressed CD45 and could be mistaken for BM-derived endometrial cells. The collective data point to a predominant role for the BM-MSC secretome, rather than differentiation capacity of MSCs, in supporting endometrial regeneration and/or repair.

In addition to the bone marrow, mesenchymal stem cells can be isolated from multiple other tissues, such as umbilical cord, adipose tissue, placenta, endometrium and menstrual blood, among others (6, 18). Although MSCs from different tissues share similar morphologic characteristics, the secretome profile of each differs significantly (18), implicating distinct functional capacities dependent on tissue source. The secretome of MSCs derived from fetal tissues such as UC has been demonstrated to have a more diverse composition that that of MSC derived from adult tissues (18). The non-invasive accessibility and distinct secretome profile of UC-MSCs highlights the need for further characterization of functional effects of UC-MSCs and their therapeutic potential in the endometrium. Our studies support a role for the UC-MSC secretome in enhancing endometrial stromal cell motility similar to that of BM-MSC, mediated in part by upregulation of CCL2 expression.

That CCL2 is a key factor mediating BM-MSC/UC-MSC-induced HESC motility is consistent with established roles for this cytokine in multiple tissues. Also known as monocyte chemoattractant protein (MCP-1), CCL2 is a pro-inflammatory chemokine known to recruit monocytes, memory T-cells, and dendritic cells (45). In cancer models, CCL2 has been implicated in stimulating tumor proliferation and metastasis by binding CCR2 and activating classic signaling cascades including MAPK/ERK signaling (46). In the endometrium, CCL2 is secreted by endometrial stromal cells, facilitates macrophage recruitment to endometriotic cells, and upregulates invasion, migration, and survival of endometrial stromal cells through paracrine and autocrine regulation, implicated in the pathogenesis of endometriosis (29, 47). CCL2 regulates the biological behavior (invasiveness and proliferation) of endometrial stromal cells in an autocrine manner by activating Akt and/or MAPK/Erk1/2 signaling pathways to promote PCNA, survivin, and MMP2 expression (29). Our data demonstrate that exposure to the BM- and UC-MSC secretomes significantly upregulated *CCL2* mRNA expression by HESCs and significantly increased T-HESC motility. Recombinant CCL2 similarly upregulated HESC migration and invasion. Since CCL2 has been identified in the MSC secretome (48), the MSC secretome is likely to upregulate HESC motility functions via both paracrine (i.e., MSC-derived CCL2 effects) as well as autocrine mechanisms.

Like CCL2, hepatocyte growth factor (HGF) has also been implicated in endometrial regeneration. HGF binds c-MET and induces activation of multiple downstream signaling pathways involved in mitogenic and motogenic functions (49). Previous studies have shown that HGF promotes proliferation, migration, and invasion of endometrial epithelial and stromal cells *in vitro* (50, 51). In the murine uterus, HGF expression is differentially modulated by estradiol and progesterone and exhibits cell type- and estrous cycle-specific changes, supporting a role for this cytokine in cyclic endometrial remodeling via autocrine/paracrine mechanisms (52). Our RNAseq studies demonstrated significant upregulation of *HGF* mRNA expression in HESCs exposed to the secretome of BM-MSC. However, qRT-PCR studies indicated only a modest but significant increase in *HGF* mRNA expression when HESC were exposed to the secretome of MSC isolated from a single BM donor, and no difference when exposed to secretome of commercially obtained BM-MSC. Similarly, exposure to the secretome of MSC isolated from one of two UC donors resulted in a modest but significant increase in *HGF* expression. These variable results, which may be attributable to inherent differences between human donors, support the need for more definitive characterization of a role for this cytokine in MSC-mediated HESC motility functions.

## Conclusions

Overall, the data presented herein demonstrate that the secretome of MSC isolated from human bone marrow and umbilical cord significantly increases HESC migratory and invasive capacities; cellular functions critical for cyclic regeneration and repair of this highly dynamic tissue. Functional effects of the MSC secretome on HESC motility appear to be mediated, in part, by upregulation of HESC CCL2 expression. Although emerging data increasingly support a potential role for mesenchymal stem cells in the treatment of defects in endometrial regeneration and function, the ability to safely and efficaciously administer these cells for therapeutic purposes remains to be established. Our data provide important implications for use of the MSC secretome as an attractive cell-free therapeutic strategy to enhance endometrial regeneration and repair after injury.

## Acknowledgements

This work was supported by the National Institutes of Health, National Heart Lung and Blood Institute (grant no. R01HL127013 to N.C.D.) and by the American Association of Obstetricians and Gynecologists Foundation (to S.S.M.).

## FIGURE LEGENDS

**Supplementary Figure 1. The secretomes of BM-MSC-3 and BM-MSC-4 increase HESC transwell migration and invasion**. HESC transwell migration and invasion through Matrigel were assessed after coculture with BM-MSCs for 24h. (**A**) Transwell migration was significantly increased by coculture with BM-MSC-3 (\*\*\*\**P*< 0.0001) and BM-MSC-4 (\*\*\**P*=0.0005) compared to control HESC. (**B**) Transwell invasion through Matrigel was significantly increased by coculture with BM-MSC-3 (\*\*\*\**P*< 0.0001), but not with BM-MSC-4 (*P*=0.06) compared to control HESC. Data are expressed as means (of N=3) ± SEM. Statistical analysis was performed with an unpaired t test followed by Welch’s correction.

