## Supplementary Tables 1 and 2 for "Bone Marrow- and Umbilical Cord-Derived Mesenchymal Stem Cell Secretome Alters Gene Expression and Upregulates Motility of Human Endometrial Stromal Cells"

**Supplementary Table 1. HESC+BM-MSC-1**

| Ingenuity Canonical Pathways | -log(p-value) | Molecules |
|---|---|---|
| DNA Methylation and Transcriptional Repression Signaling | 12.10 | H4C1,H4C11,H4C13,H4C2,H4C3,H4C4,H4C6,H4C9 |
| Sirtuin Signaling Pathway | 11.70 | H1-1,H1-2,H1-3,H1-4,H1-5,H3C3,H4C11,MT-ATP6,MT-CYB,MT-ND1,MT-ND2,MT-ND4,MT-ND4L,MT-ND5,MT-ND6 |
| Transcriptional Regulatory Network in Embryonic Stem Cells | 10.50 | H4C1,H4C11,H4C13,H4C2,H4C3,H4C4,H4C6,H4C9 |
| NAD Signaling Pathway | 9.00 | H1-1,H1-2,H1-3,H1-4,H1-5,H2BC10,H2BC17,H2BC8,H2BU1,HGF |
| NER (Nucleotide Excision Repair, Enhanced Pathway) | 8.09 | H4C1,H4C11,H4C13,H4C2,H4C3,H4C4,H4C6,H4C9 |
| Granzyme A Signaling | 8.09 | H1-1,H1-2,H1-3,H1-4,H1-5 |
| Glucocorticoid Receptor Signaling | 6.39 | CCL2,H3C3,KRT18,MT-ATP6,MT-CYB,MT-ND1,MT-ND2,MT-ND4,MT-ND4L,MT-ND5,MT-ND6,NR3C2,PLAU |
| Oxidative Phosphorylation | 6.37 | MT-ATP6,MT-CYB,MT-ND1,MT-ND2,MT-ND4,MT-ND4L,MT-ND5 |
| Mitochondrial Dysfunction | 6.19 | MT-ATP6,MT-CYB,MT-ND1,MT-ND2,MT-ND4,MT-ND4L,MT-ND5,MT-ND6 |
| Ferroptosis Signaling Pathway | 4.42 | H2AC18/H2AC19,H2AC21,H2BC10,H2BC17,H2BC8,H2BU1 |
| Estrogen Receptor Signaling | 3.72 | MT-ATP6,MT-CYB,MT-ND1,MT-ND2,MT-ND4,MT-ND4L,MT-ND5,MT-ND6 |
| Protein Kinase A Signaling | 2.93 | AKAP9,H1-1,H1-2,H1-3,H1-4,H1-5,H3C3 |
| Coagulation System | 2.10 | BDKRB2,PLAU |
| Kinetochore Metaphase Signaling Pathway | 2.05 | CENPE,H2AC18/H2AC19,H2AC21 |
| Tumor Microenvironment Pathway | 1.45 | CCL2,HGF,PLAU |
| Renal Cell Carcinoma Signaling | 1.40 | HGF,VHL |

**Supplementary Table 2. HESC+BM-MSC-2**

| Ingenuity Canonical Pathways | -log(p-value) | Molecules |
|---|---|---|
| Role of Hypercytokinemia/hyperchemokinemia in the Pathogenesis of Influenza | 8.74 | CASP1,CCL2,DDX58,IRF7,ISG15,MX1, OAS2,OAS3 |
| Interferon Signaling | 8.27 | IFI35,IFI6,IFIT1,IFITM1,ISG15,MX1 |
| Coronavirus Pathogenesis Pathway | 6.98 | BST2,CASP1,CCL2,DDX58,FOS,IRF7,OAS2, OAS3,PTGS2 |
| Tumor Microenvironment Pathway | 6.12 | CCL2,FGF9,FOS,FOXO1,HGF,PDGFB,PLAU, PTGS2 |

| | | |
|---|---|---|
| Role of Pattern Recognition Receptors in Recognition of Bacteria and Viruses | 5.57 | CASP1,DDX58,IFIH1,IRF7,OAS2,OAS3, TNFSF4 |
| Activation of IRF by Cytosolic Pattern Recognition Receptors | 5.25 | DDX58,DHX58,IFIH1,IRF7,ISG15 |
| Regulation Of The Epithelial Mesenchymal Transition By Growth Factors Pathway | 4.98 | EGR1,FGF9,FOS,FOXO1,HGF, PDGFB,TNFSF4 |
| Role of RIG1-like Receptors in Antiviral Innate Immunity | 4.50 | DDX58,DHX58,IFIH1,IRF7 |
| Role of PKR in Interferon Induction and Antiviral Response | 3.69 | CASP1,DDX58,FOS,IFIH1,PDGFB |
| Pulmonary Fibrosis Idiopathic Signaling Pathway | 3.52 | EGR1,FGF9,FOS,FOXO1,PDGFB, PLAU,WNT2 |
| Neuroinflammation Signaling Pathway | 3.48 | CASP1,CCL2,FOS,GDNF,GRIA1,IRF7, PTGS2 |
| IL-17A Signaling in Fibroblasts | 3.36 | CCL2,CEBPD,FOS |
| Endocannabinoid Neuronal Synapse Pathway | 3.34 | CACNA1H,GRIA1,GRIA4,PLCH1,PTGS2 |
| Neurovascular Coupling Signaling Pathway | 3.34 | BDKRB2,CACNA1H,GRIA1,GRIA4, KCNMB4,PTGS2 |
| CREB Signaling in Neurons | 3.17 | ADRA2C,BDKRB2,CACNA1H,EDNRB, GRIA1,GRIA4,HGF,PDGFB,PLCH1 |
| Neuropathic Pain Signaling In Dorsal Horn Neurons | 3.11 | FOS,GRIA1,GRIA4,PLCH1 |
| Endothelin-1 Signaling | 2.88 | CASP1,EDNRB,FOS,PLCH1,PTGS2 |
| Amyotrophic Lateral Sclerosis Signaling | 2.84 | CASP1,GDNF,GRIA1,GRIA4 |
| Prostanoid Biosynthesis | 2.76 | CYP2S1,PTGS2 |
| Role of Macrophages, Fibroblasts and Endothelial Cells in Rheumatoid Arthritis | 2.70 | CCL2,CEBPD,FOS,PDGFB,PLCH1, WNT2 |
| Adipogenesis pathway | 2.70 | CEBPD,FOXO1,RBP1,RUNX1T1 |
| Thyroid Cancer Signaling | 2.40 | FOS,FOXO1,GDNF |
| Renal Cell Carcinoma Signaling | 2.39 | FOS,HGF,PDGFB |
| Glioblastoma Multiforme Signaling | 2.30 | FOXO1,PDGFB,PLCH1,WNT2 |
| TR/RXR Activation | 2.26 | AKR1C1/AKR1C2,BCL3,DIO2 |
| ERBB Signaling | 2.22 | EREG,FOS,FOXO1 |
| Cardiac Hypertrophy Signaling (Enhanced) | 2.22 | ADRA2C,EDNRB,FGF9,PLCH1,PTGS2, TNFSF4,WNT2 |
| Pyroptosis Signaling Pathway | 2.20 | CASP1,GBP2,GBP4 |

| | | |
|---|---|---|
| IL-17 Signaling | 2.19 | CCL2,FOS,PTGS2,TNFSF4 |
| UVA-Induced MAPK Signaling | 2.16 | FOS,PARP9,PLCH1 |
| Regulation of the Epithelial-Mesenchymal Transition Pathway | 2.14 | EGR1,FGF9,HGF,WNT2 |
| Hepatic Fibrosis / Hepatic Stellate Cell Activation | 2.12 | CCL2,EDNRB,HGF,PDGFB |
| Oxytocin In Brain Signaling Pathway | 2.09 | CACNA1H,CASP1,PTGS2,RGS2 |
| Coagulation System | 2.08 | BDKRB2,PLAU |
| Synaptic Long Term Depression | 2.07 | CACNA1H,GRIA1,GRIA4,PLCH1 |
| PPAR Signaling | 2.07 | FOS,PDGFB,PTGS2 |
| Airway Pathology in Chronic Obstructive Pulmonary Disease | 1.95 | CCL2,FGF9,TNFSF4 |
| Autophagy | 1.92 | FOS,FOXO1,HGF,PDGFB |
| Sphingosine-1-phosphate Signaling | 1.90 | CASP1,PDGFB,PLCH1 |
| Retinoate Biosynthesis I | 1.86 | AKR1C1/AKR1C2,RBP1 |
| 14-3-3-mediated Signaling | 1.85 | FOS,FOXO1,PLCH1 |
| MIF Regulation of Innate Immunity | 1.84 | FOS,PTGS2 |
| Osteoarthritis Pathway | 1.82 | CASP1,DDIT4,PTGS2,RARRES2 |
| HGF Signaling | 1.80 | FOS,HGF,PTGS2 |
| Synaptic Long Term Potentiation | 1.76 | GRIA1,GRIA4,PLCH1 |
| CSDE1 Signaling Pathway | 1.69 | CCL2,FOS |
| Estrogen Biosynthesis | 1.69 | AKR1C1/AKR1C2,CYP2S1 |
| White Adipose Tissue Browning Pathway | 1.66 | CACNA1H,DIO2,RUNX1T1 |
| Role of IL-17A in Arthritis | 1.65 | CCL2,PTGS2 |
| Thyroid Hormone Metabolism II (via Conjugation and/or Degradation) | 1.62 | CSGALNACT1,DIO2 |
| Circadian Rhythm Signaling | 1.61 | CACNA1H,GRIA1,GRIA4,PLCH1 |
| Ovarian Cancer Signaling | 1.61 | FGF9,PTGS2,WNT2 |
| Corticotropin Releasing Hormone Signaling | 1.60 | CACNA1H,FOS,PTGS2 |
| Nicotine Degradation III | 1.60 | CSGALNACT1,CYP2S1 |
| Cancer Drug Resistance By Drug Efflux | 1.57 | FOXO1,PTGS2 |
| Hepatic Fibrosis Signaling Pathway | 1.57 | CCL2,FOS,FOXO1,PDGFB,WNT2 |

| | | |
|---|---|---|
| HMGB1 Signaling | 1.56 | CCL2,FOS,TNFSF4 |
| CD40 Signaling | 1.55 | FOS,PTGS2 |
| NAD Signaling Pathway | 1.54 | HGF,PARP9,PDGFB |
| Human Embryonic Stem Cell Pluripotency | 1.53 | FOXO1,PDGFB,WNT2 |
| Aldosterone Signaling in Epithelial Cells | 1.51 | KCNMB4,NR3C2,PLCH1 |
| Retinoate Biosynthesis II | 1.51 | RBP1 |
| Melatonin Degradation I | 1.48 | CSGALNACT1,CYP2S1 |
| Glutamate Receptor Signaling | 1.48 | GRIA1,GRIA4 |
| Nicotine Degradation II | 1.45 | CSGALNACT1,CYP2S1 |
| Macropinocytosis Signaling | 1.45 | HGF,PDGFB |
| TREM1 Signaling | 1.43 | CASP1,CCL2 |
| IL-3 Signaling | 1.42 | FOS,FOXO1 |
| GDNF Family Ligand-Receptor Interactions | 1.42 | FOS,GDNF |
| Leptin Signaling in Obesity | 1.41 | FOXO1,PLCH1 |
| IL-7 Signaling Pathway | 1.41 | FOXO1,HGF |
| Role of MAPK Signaling in Inhibiting the Pathogenesis of Influenza | 1.41 | CCL2,PTGS2 |
| Neurotrophin/TRK Signaling | 1.38 | FOS,SPRY1 |
| GNRH Signaling | 1.37 | CACNA1H,EGR1,FOS |
| Chemokine Signaling | 1.37 | CCL2,FOS |
| PPARŒ±/RXRŒ± Activation | 1.36 | BCL3,HELZ2,PLCH1 |
| Estrogen-Dependent Breast Cancer Signaling | 1.36 | AKR1C1/AKR1C2,FOS |
| Role of MAPK Signaling in the Pathogenesis of Influenza | 1.36 | CCL2,PTGS2 |
