## Supplementary figures and images for "Bone Marrow- and Umbilical Cord-Derived Mesenchymal Stem Cell Secretome Alters Gene Expression and Upregulates Motility of Human Endometrial Stromal Cells"

### Supplemental Figure 1

#### Slide 1
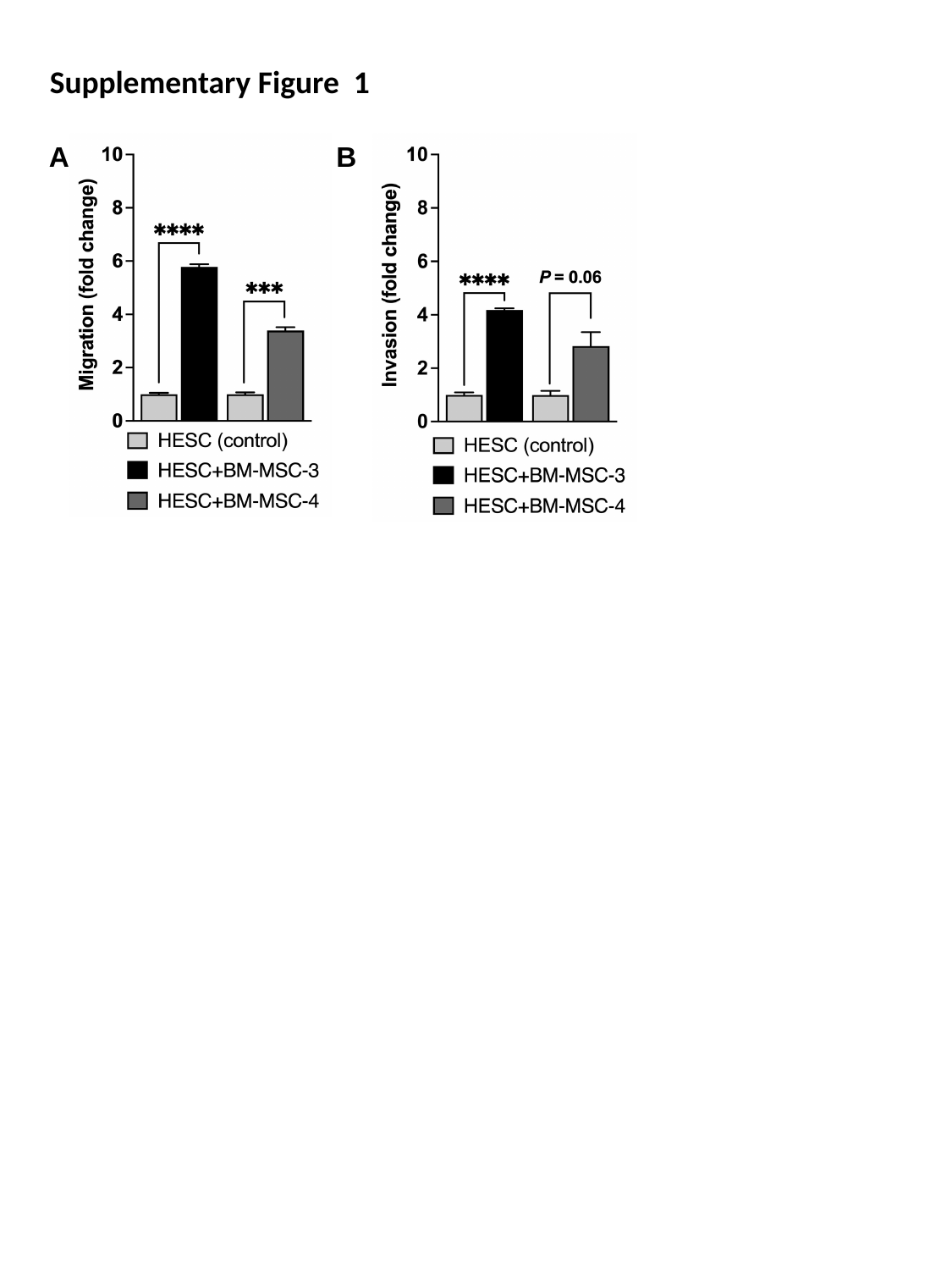

Supplementary Figure 1
A
B
